# Improving model-based fNIRS analysis using mesh-based anatomical and light-transport models

**DOI:** 10.1101/2020.02.07.939447

**Authors:** Anh Phong Tran, Shijie Yan, Qianqian Fang

**Affiliations:** Northeastern University, Dept. of Chemical Engineering, Boston, Massachusetts, United States; Northeastern University, Dept. of Electrical and Computer Engineering, Boston, Massachusetts, United States; Northeastern University, Dept. of Bioengineering, Boston, Massachusetts, United States

**Keywords:** functional near-infrared spectroscopy, tetrahedral mesh generation, brain atlas, Monte Carlo method

## Abstract

**Significance:** Functional near-infrared spectroscopy (fNIRS) has become an important research tool in studying human brains. Accurate quantification of brain activities via fNIRS relies upon solving computational models that simulate the transport of photons through complex anatomy.

**Aim:** We aim to highlight the importance of accurate anatomical modeling in the context of fNIRS, and propose a robust method for creating high-quality brain/full-head tetrahedral mesh models for neuroimaging analysis.

**Approach:** We have developed a surface-based brain meshing pipeline that can produce significantly better brain mesh models compared to conventional meshing techniques. It can convert segmented volumetric brain scans into multi-layered surfaces and tetrahedral mesh models, with typical processing times of only a few minutes and broad utilities, such as in Monte Carlo or finite-element based photon simulations for fNIRS studies.

**Results:** A variety of high quality brain mesh models have been successfully generated by processing publicly available brain atlases. In addition, we compare 3 brain anatomical models - the voxel-based brain segmentation, tetrahedral brain mesh and layered-slab brain model, and demonstrate noticeable discrepancies in brain partial-pathlengths when using approximated brain anatomies, ranging between −1.5-23% with the voxelated brain and 36-166% with the layered-slab brain.

**Conclusion:** The generation and utility of high-quality brain meshes can lead to more accurate brain quantification in fNIRS studies. Our open-source meshing toolboxes “Brain2Mesh” and “Iso2Mesh” are freely available at http://mcx.space/brain2mesh.

## 1 Introduction

Functional near-infrared spectroscopy (fNIRS) has played an increasingly important role in functional neuroimaging.^1^ Using light in the red and near-infrared range, the haemodynamic response of the brain is probed through careful placement of sources and detectors on the scalp surface at multiple wavelengths. Relative changes in oxygenated (HbO_2_) and deoxygenated hemoglobin (HbR) concentrations, as a result of neural activities, lead to variations in light intensities at the detectors that are used to infer the locations of these activities. The accuracy of this inference depends greatly not only on an accurate representation of the complex human brain anatomy, but also the surrounding tissues that affect the migrations of photons from the sources to the detectors. Using the anatomical scans of the patients head or a resembling atlas, Monte Carlo (MC) simulations are often used in conjunction with tissue optical properties to approximate the photon path-lengths that are used to reconstruct the changes in HbR and HbO_2_. While much simplified brain models, such as planar^2, 3^ or spherical layers,^4^ as well as approximated photon propagation models, such as the diffusion approximation,^5^ have been widely utilized by the research community, their limitations are recognized by a number of studies.^3, 6^ In addition, modeling brain anatomy accurately also plays important roles in other quantitative neuroimaging modalities, such as electroencephalography (EEG)^7^ and magnetoencephalography (MEG).^8^

Whereas a voxelated brain representation has been dominantly used for acquiring and storing a three-dimensional (3-D) neuroanatomical volume, the terraced boundary shape in a voxelated space has difficulty in representing a smooth and curved boundary that typically delineates human tissues, resulting in a loss of accuracy, especially when modeling complex cortical surfaces with limited resolution. In addition, the uniform grid structure of the voxel space also demands a large number of cells in order to store brain anatomy without losing spatial details; this can cause prohibitive memory allocation and runtime in applications where solving sophisticated numerical models is necessary. Another approach – octree – uses nested voxel refinement near curved boundaries. This improves memory efficiency significantly, but still suffers from terraced mesh boundaries.^9–11^

Mesh-based brain/head models made of triangular surfaces or tetrahedral elements have advantages in both improved boundary accuracy and high flexibility compared to voxelated domains. Mesh-based models are not only the most common choice in computer graphics and 3-D visualization of brain structures, they are also the primary format for finite element analysis (FEA) and image reconstructions in many neuroimaging studies.^12, 13^ In fNIRS, tetrahedral meshes have been reported in several studies to model light propagation and recover brain hemodynamic activation using the finite-element^14, 15^ or mesh-based Monte Carlo (MMC) method.^16, 17^ Despite its importance for quantitative fNIRS analysis, the available mesh-based brain models remain limited^6, 18, 19^ in part due to the difficulties in generating accurate brain tetrahedral meshes.

The importance of creating high-quality brain mesh models is not limited to fNIRS. EEG relies on surface and volumetric head/brain meshes to quantitatively estimate the brain cortical activities.^20^ The effects of transcranial magnetic stimulation (TMS) and transcranial direct current stimulation (tDCS) can also be simulated on realistic mesh models to evaluate brain damages^13, 21^ or by measuring tDCS effects on major brain disorders.^22^ In addition, FEA of brain tissue deformation using mesh models can assist neurosurgeons in the study of traumatic brain injuries or surgical planning.^23^

On the other hand, while it has been generally agreed that mesh-based light transport models are more accurate than those using a voxel-based domain, there has not been a systematic study to investigate such difference and its impact to fNIRS. The shape differences between a voxelated and a mesh-based boundary not only influence how photons get absorbed by tissue, thus altering fluence distributions, but also impact greatly on photon reflection/transmission characteristics near a tissue/air boundary. Such error could also be amplified with the presence of low-scattering/low-absorption tissue, such as cerebrospinal fluid (CSF) in the brain, and result in inaccurate estimations in fNIRS. To quantify the impact of brain anatomical models in light modeling using the state-of-the-art voxel and mesh-based MC simulators could be greatly beneficial for the community to design more efficient study protocols and instruments.

Despite the broad awareness of anatomical model differences, fNIRS studies utilizing mesh brain models are quite limited, largely due to the challenges to create brain meshes and lack of publicly available meshing tools. A large portion of these studies rely on previously created meshes^24^ or using general-purpose tetrahedral mesh generators that are not optimized for meshing the brain. For example, a voxel conforming mesh generation approach,^25, 26^ the marching cubes algorithm^27, 28^ and the “Cleaver” software,^29^ can achieve good surface accuracy, but at the cost of highly dense elements near the boundaries due to the octree-like refinement. A general-purpose 3-D mesh generation pipeline was proposed alongside with an open-source meshing software BioMesh3D.^30^ This approach makes use of physics-based optimizations to obtain a high-quality multi-material feature-preserving tetrahedral mesh models, but at the expense of lengthy run-time, on the order of several hours. In another Delaunay-based meshing pipeline provided in the Computational Geometry Algorithms Library (CGAL), also supported in Iso2Mesh^31^ in 2009 and NIRFAST in 2013,^32^ the tetrahedral mesh is generated from a random point-set that is iteratively refined.^33, 34^ This procedure is relatively fast, parallelizable and robust; however, non-smooth boundaries are often observed between tissue regions (example shown below). Commercially available tools, such as Mimics (Materialise, Leuven, Belgium), and ScanIP (Simpleware, Exeter, UK), offer integrated interfaces for image segmentation and mesh generation, but often require manual mesh editing; a streamlined high-quality mesh processing pipeline from neuroanatomical scans remains challenging.

Meshing tools specifically optimized for brain mesh generation are very limited. In 2010, we reported a surface-based brain meshing approach.^16^ In 2011, a similar approach was reported,^35^ in which an automated meshing pipeline “mri2mesh” was reported, incorporating FreeSurfer surface models and the scalp, skull, and CSF segmentations from FSL with the assistance of a surface decoupling step.^36^ Although mri2mesh yielded smooth boundaries and high-quality tetrahedral elements, the reported meshing times are on the order of 3-4 hours, in addition to the time for segmentation. Moreover, mri2mesh can only process FreeSurfer and FSL outputs, limiting its integration with other segmentation tools.

In this work, we address two challenges in model-based neuroimaging analysis. First, we report a fully automated, surface-based brain/full head 3-D meshing pipeline “Brain2Mesh” – a specialized wrapper built upon our widely-adopted mesh generation toolbox, “Iso2Mesh”, dedicated towards high-quality brain mesh generation. A major difference separating this work from the more conventional CGAL-based volumetric meshing approach^32, 33^ is the use of surface-based meshing workflow. This allows it to produce brain mesh models with significantly higher quality. It is also much more flexible, processing data conveniently from multi-label (discrete) or probabilistic (continuous) segmentations and surface models of the brain. Secondly, this work quantitatively demonstrates that by utilizing an accurate mesh-based brain representation, one can potentially improve the accuracy in fNIRS data analysis. We analyze the errors in both fluence and photon partial-pathlengths in the MC simulation outputs comparing between a layered-slab, a voxel-based and a mesh-based brain model.

In the remainder of this paper, we first detail the brain mesh generation pipeline that we have developed to create high-quality brain mesh models. In the subsequent Results and Discussion sections, we show a varieties of examples of the brain/full head meshes created using different types of neuroanatomical data inputs, including an open-source brain mesh libraries created based on the recently published Neurodevelopmental MRI Database.^37^ In addition, we also perform mesh-based MC simulation using the produced mesh-model and compare the results with those from voxelated and layered-slab brain models. We highlight the modeling errors of using a voxelated model in both fluence and partial-pathlength in comparison with a mesh-based brain.

## 2 Material and methods

### 2.1 High-quality 3-D brain mesh generation pipeline

#### 2.1.1 Brain segmentation

The brain anatomical modeling pipeline reported in this work starts from a pre-segmented brain. Here, we want to particularly highlight that brain segmentation is outside the scope of this paper. As noted below, there is an array of dedicated brain segmentation tools, extensively developed and validated over the past several decades. Advanced statistical and template-based segmentation methods have been investigated and rigorously implemented in these tools. It is not our interests to develop a new segmentation method, but to convert these pre-segmented anatomies into accurate meshes for subsequent computational modeling.

A diagram summarizing the common pathways in segmenting a neuroanatomical scan using popular neuroimaging analysis tools is shown in Fig. 1. In most cases, a tissue probability or multi-label volume is obtained for the white-matter (WM), gray-matter (GM) and CSF. Some segmentation tools, such as Statistical Parametric Mapping (SPM), allow the use of a matching T2-weighted MRI to improve the CSF segmentation. Utilities such as the FSL brain extraction tool and SPM can provide additional information on the scalp and skull. In this work, we primarily focus on 6 tissue types – WM, GM, CSF, skull, scalp and air cavities. Additional classes of tissue (e.g. dura, vessels, fatty tissues, skin, muscle) are also available in some segmentation outputs, which can be incorporated to our mesh generation pipeline. However, one should be aware that adding additional segmentations may result in increased node numbers and surface complexity, including disconnected surface components.

**Fig 1.**
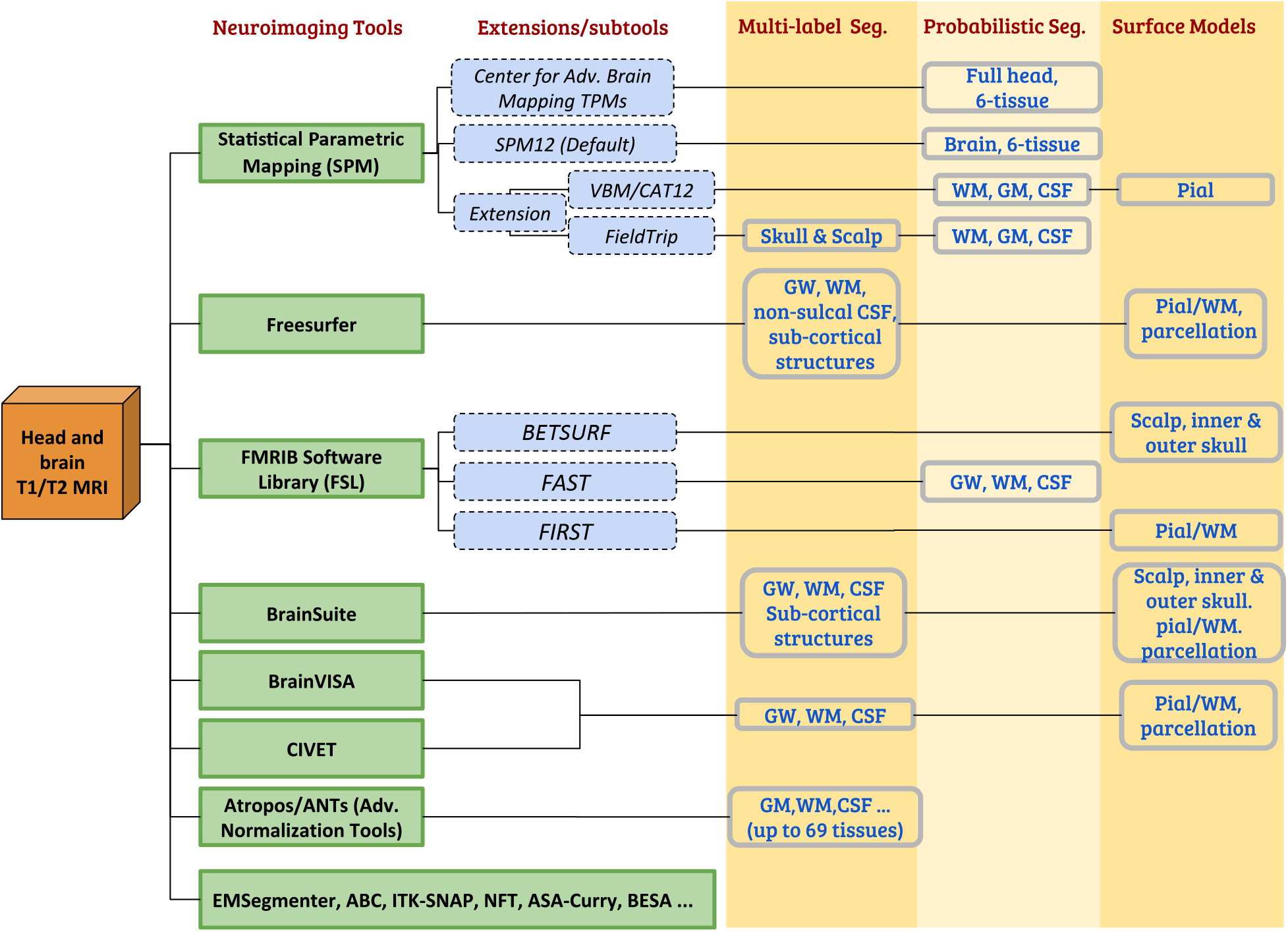
Segmentation pathways from anatomical head and brain MRI scans. The common neuroimaging tools/extensions (left) and the corresponding outputs (right, shaded) are listed.

#### 2.1.2 Segmentation pre-preprocessing

The segmented brain or full-head volume, represented by either a multi-label 3-D mask or a set of probability maps (in floating-point 0-1 values), are preprocessed to ensure a layered tissue model i.e., the WM, GM, CSF, bone and scalp are incrementally enclosed by or sharing a common boundary with the later tissue layers – the scalp surface is the outermost layer and the WM is the inner-most layer. In previous publications, two adjacent layers must be separated by a non-zero gap.^35, 38^ In this work, we have overcome this limitation by initially inserting a small gap between successive layers and then applying a post-processing step to recover the merged boundaries. Moreover, we also consider air cavities, which can be located between two tissue surfaces, for example, inside or outside skull surfaces. In Fig. 2, the workflow to create different brain tissue boundaries is outlined.

**Fig 2.**
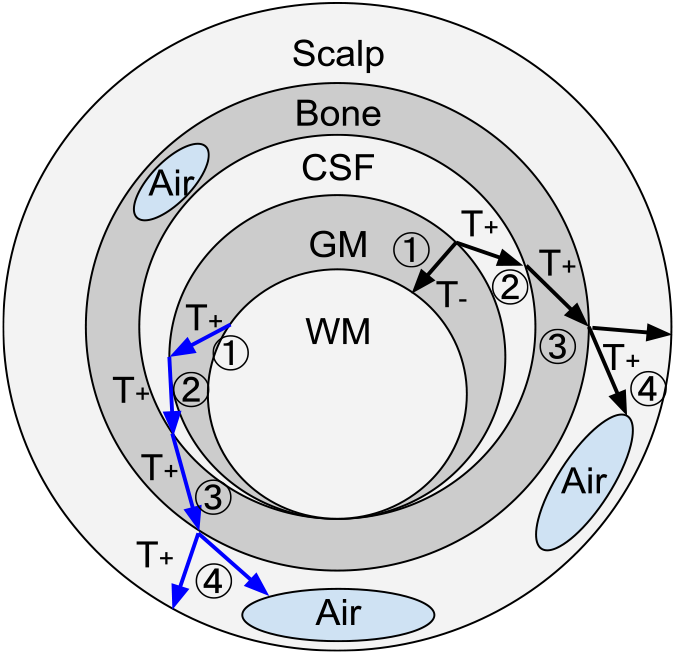
Illustration of the layered tissue model and the segmentation preprocessing workflow. Multiple air cavities are allowed. An arrow represents a thinning (*T*_−_) or thickening (*T*_+_) operation between two adjacent regions. Two sample pathways are indicated, shown by black and blue arrows, respectively. The circled numbers indicate the processing order. The gaps inserted between layers can be removed in the post-processing step to recover shared boundaries (such as the CSF/GM/WM surfaces here).

**Fig 3.**
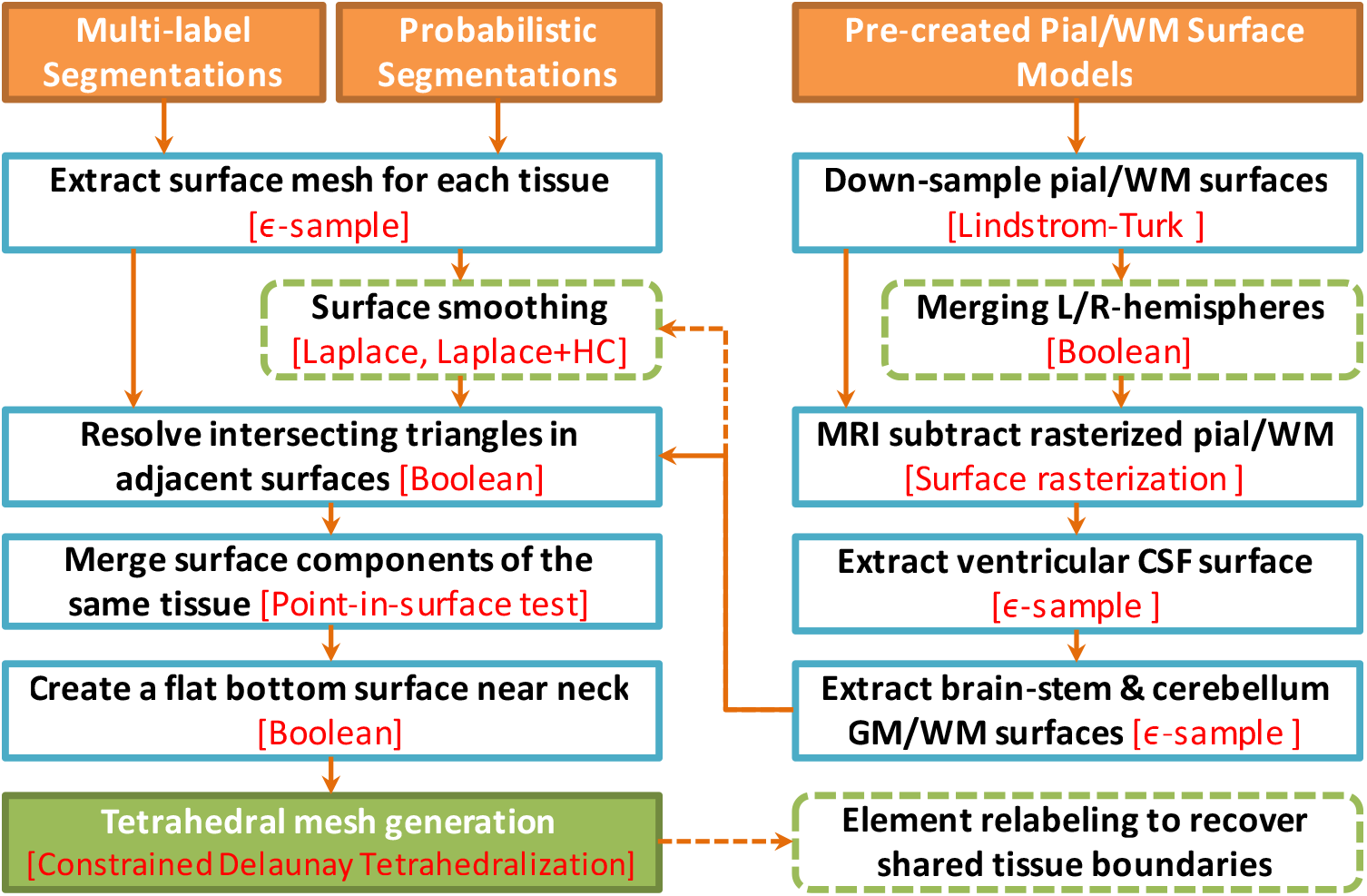
Processing steps for a surface-based mesh generation workflow. The left-side shows the steps for processing tissue probability maps and multi-label volumes, and the right-side shows additional steps to incorporate pre-created pial and white matter surfaces. The specific algorithm used in each step is indicated in red, while dashed boxes and arrows indicate optional processing steps.

To avoid intersecting triangles in the generated multi-layered surface model, we first insert a small gap between adjacent brain tissue layers (only in the regions where tissues have shared boundaries).

This is achieved using either a 3-D image “thickening” or “thinning” operator in the segmented volume. In the case of a thickening operator (*T*_+_), the outer layer tissue segmentation (*P_out_*) is modified by

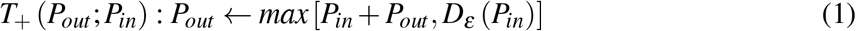

where *P_in_* and *P_out_* represent the inner and outer tissue probabilistic segmentation, respectively. *D* denotes a “max-filter”, i.e. a volumetric dilation operator defined by replacing each voxel with the maximum value in a cubic neighboring region with a half-edge length of *ε*, i.e.

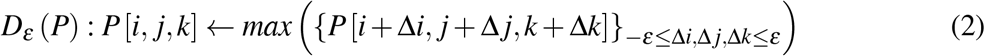

Similarly, a “thinning” operator (*T*_−_) is defined by

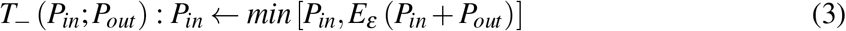

where *E_ε_* (*P_out_*) is an “erosion” operator of width *ε*, defined by a “min-filter” as

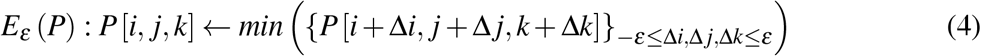

The *T*_−_ operator effectively shrinks the inner layer mask at the intersecting regions with the outer layer. We note here that the above *T*_+_ and *T*_−_ operators work for both probabilistic and binary segmentations. We also highlight that the above operators only alter tissue segmentations in the regions where the inner tissue boundaries merge or intersect with the outer boundaries. Such regions only account for a small fraction of the brain tissue boundaries generated from realistic data. An optional post-processing step is applied to “relabel” the elements in the expanded regions to recover the shared tissue boundaries (see below).

#### 2.1.3 Tissue surface extraction and surface-mesh processing

If the input data are given in the form of a multi-labeled volume or tissue probability maps (see Fig. 1), the next step of the mesh generation is to create a triangular surface mesh for each tissue layer. This is achieved using the “ *ε*-sample” algorithm using the CGAL Surface Mesh Generation library.^39^ For each extracted tissue surface, independent mesh quality and density criteria can be defined. In general, a surface mesh extracted from a probabilistic segmentation (grayscale) is smoother than one derived from a binary segmentation. In addition, we also provide 3 surface smoothing algorithms, including the Laplacian, Laplacian+HC and low-pass filters.^40^ Moreover, a surface boolean operation using on a customized Cork 3-D surface Boolean library^41^ is used to avoid surface intersections.

If the segmentation tool directly outputs the surface meshes of GM and WM, such as FreeSurfer, additional surface preprocessing is often required. For example, the FreeSurfer GM/WM surfaces are typically very dense. A surface simplification algorithm using the Lindstrom-Turk algorithm is applied^42, 43^ to decimate the surface nodes. In another example, if the pial and white matter surfaces contain a separate surface for each brain hemisphere, a “merge” operation is applied to combine them into one closed surface. Additionally, the FreeSurfer and FSL pial/WM surfaces do not cover the cerebellum and brainstem regions. To add those two anatomical regions, we first rasterize the pial/WM surfaces and then subtract those from the GW/WM probabilistic segmentations. From the subtracted probability maps, we extract the brainstem and cerebellum white and gray matter surfaces, and subsequently merge these meshes with the cerebral GM/WM surfaces.

#### 2.1.4 Volumetric mesh generation and post-processing

With the above derived combined multi-layer head surface model, a full head tetrahedral mesh can be finally generated using a constrained Delaunay tetrahedralization (CDT) algorithm, achieved using an open-source meshing utility TetGen.^44^ The output mesh quality and density are fully controlled by a set of user-defined meshing criteria. An unnormalized radius-edge ratio (*q*) lower bound (see below) can be specified to control the overall quality of the tetrahedral elements. In addition, one can set an upper-bound for the tetrahedral element volume globally for the entire mesh, or for a particular tissue label. Spatially varying mesh density is also supported via user-defined “sizing-field”.^44^ After tessellation, each enclosed region in a multi-layered brain surface model is filled by tetrahedral elements, and is assigned with a unique region label to distinguish different tissue types.

If the small gaps inserted by the aforementioned thickening and thinning operations are not desired, an optional “relabeling” step is performed to recover the originally merged tissue boundaries. To do this, we use the centroids of the tetrahedral mesh elements in each unique region to determine the innermost surface that encloses this region volumes, based on which we can re-tag these elements using the correct tissue labels.

### 2.2 Assessing impact of anatomical models in light transport modeling

The ability to generate high-quality brain mesh models alongside with in-depth understandings to the state-of-the-art voxel and mesh-based MC photon transport modeling tools allow us to investigate the impact of brain anatomical models on fNIRS data analysis. Here, we are interested in quantitatively comparing various brain anatomical models in terms of light transport modeling, and quantify their differences in the optical parameters essential to fNIRS measurements. The three brain anatomical models that we are evaluating include 1) mesh-based brain models,^6, 16^ 2) voxel-based brain segmentations,^45^ and 3) simple layered-slab brain models often found in literature.^3, 46^ We apply our widely adopted and cross-validated MC light transport simulators – Mesh-based MC (MMC)^16, 17^ for mesh- and layered-slab brain simulations, and Monte Carlo eXtreme (MCX)^31, 47^ for voxel-based brain simulations. Furthermore, a number of publications applied the diffusion approximation (DA) to fNIRS modeling by utilizing an approximated scattering coefficient for CSF,^5, 48^ as it has very low scattering. It is beneficial to the community to understand the errors caused by such model approximation.

To ensure that the mesh model is “equivalent” to the original segmentation, we calculate the volumetric ratios, denoted as *V_rel_*, between the enclosed volume of the tissue boundary over those derived from the corresponding segmentation for a given tissue. A *V_rel_* value close to 1 suggests excellent volume conservation. From all MC simulations, we compute important light transport parameters, such as the average partial-pathlengths^49^ in the GM/WM regions (*PPL_B_* in mm), the fraction of the partial-pathlengths in the brain (*R_B_*) as well as the optical fluence spatial distributions. In addition, we also compute the percentage fractions of the total energy deposition in the brain regions using both the mesh and voxel-based simulations. Such parameter is strongly relevant to photobiomodulation (PBM) applications.^50^

## 3 Results and Discussion

In the below subsections, we first showcase the robustness and flexibility of our aforementioned brain meshing pipeline by processing a wide range of complex brain anatomical scans. Various meshing pathways of our meshing pipeline are validated by using publicly available brain atlas datasets, including the Neurodevelopmental MRI database.^37, 51^ In addition, we use a sample full-head mesh generated from the MRI database and report their differences in key optical parameters by performing 3-D mesh-, voxel- and layered-domain Monte Carlo transport simulations at a range of source-detector separations. All computational times were benchmarked on an Intel i7-8700K processor using a single thread.

### 3.1 High quality tetrahedral meshes of human head and brain models

In Fig. 4, a sample full-head mesh model is generated from an SPM segmentation with tissue priors from the Laboratory for Research in Neuroimaging (LREN).^52^ The generated mesh contains 181,026 nodes and 1,060,051 tetrahedral elements. A T1-weighted MRI scan of an average head for the 40-44 years old age group from the University of South Carolina (USC) Neurodevelopmental MRI database (in the following sections, we will use “USC age-group” to refer to an atlas from this database, for example, the atlas used in this example is USC 40-44) was used as the input. The segmentation yields 5 tissue classes that are used in the mesh generation: WM, GM, CSF, bone, and scalp.

**Fig 4.**
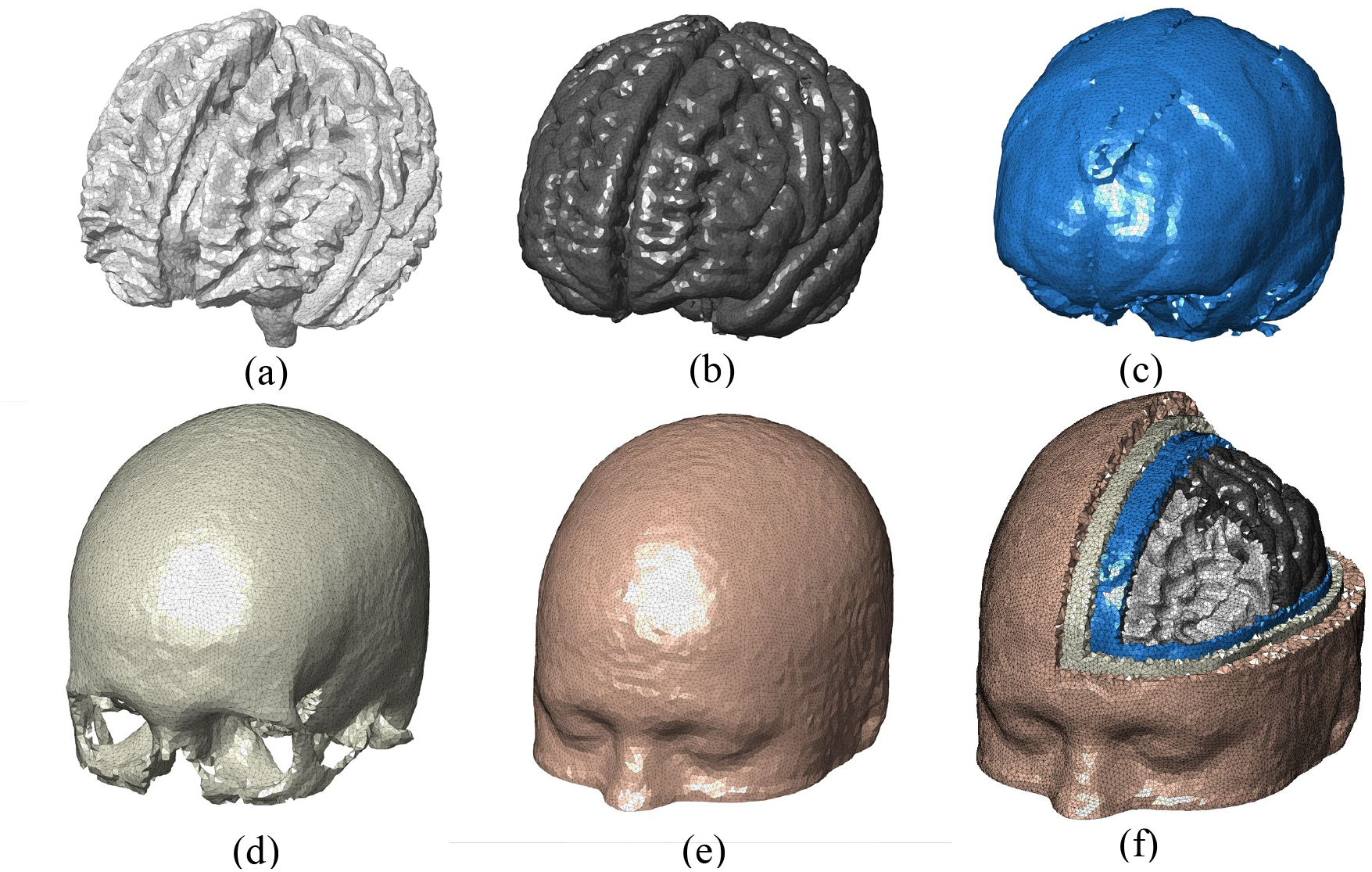
A 5-layer full head tetrahedral mesh derived from an atlas head of the USC 40-44 atlas. It contains (a) WM, (b) GM, (c) CSF, (d) bone, and (e) scalp layers. A cross-cut view of the tetrahedral mesh is shown in (f).

The mesh density for each tissue surface and volumetric region is fully customizable by setting the following three parameters:

- *R_max_*: the maximum radius of the Delaunay sphere^39^ (in voxel unit) that bounds each of the triangles in a given surface mesh. For example, setting *R_max_* = 2 requires that the circumscribed sphere of each surface triangle in the generated mesh must have a radius less than 2 (voxel-length). A smaller *R_max_* value is generally needed when meshing objects with sharp features.
- *V_max_*: the maximum tetrahedral element volume^44^ (in cubic voxel-length unit). For example, setting *V_max_* to 4 means no tetrahedral element in the generated mesh can exceed 4 cubic voxels in volume.
- *q*: the lower-bound of the radius-to-edge ratio^44^ (measuring mesh quality), defined by *q* = min({*R_i_/L_i_*}_*i*=1,2,…_) where *R_i_* is the radius of the circumsphere of the *i*-th tetrahedron, and *L_i_* is the shortest-edge length of that tetrahedron.

Each surface of the brain tissue layer is extracted using a layer-specific *R_max_* value: *R_max_* = 1.7 mm for pial and WM, 2 mm for CSF, 2.5 mm for skull and 3.5 mm for scalp. The *V_max_* value was defined as 30 mm^3^, and *q* was set to 1.414. The probability threshold for surface extraction was set at 0.5 for each of the tissues. The entire mesh takes 53.47 seconds to generate using a single CPU thread. The *V_rel_* values computed from the generated mesh layers for WM, GM, CSF, skull and scalp are 0.9924, 0.9989, 0.9921, 1.0088 and 1.0037, respectively. The excellent match of the volumes is a strong indication that our meshing pipeline preserves the tissue shapes accurately.

The tissue surfaces shown in Fig. 4 are visually smooth with an average Joe-Liu quality metric^53^ of 0.74, indicating excellent element shape quality. Overall, no degenerated element is found. The element volumes in Fig. 4 are well distributed (not shown) with only a small portion of relatively small elements. The significant improvement in mesh quality is demonstrated in the side-by-side comparison shown in Fig. 5. Here we compare the full head meshes created using the conventional CGAL-based direct meshing approach^32^ and the results from our surface-based meshing pipeline. From (a-c), there are several notable limitations from the conventional approach, namely 1) the tissue boundaries are not smooth (also evident from Fig. 2 in ^32^), 2) it has difficulty processing thin-layered tissue such as CSF (see Fig. 5b), and 3) it can produce small isolated “islands” (see Fig. 5c) due to the noise present in the volumetric image. In comparison, our surface-based meshing pipeline produces smooth and well-shaped surfaces with correct topological order. We also want to highlight that the CGAL-generated mesh contains over 268,000 nodes, while the mesh from our approach has only 158,211 nodes.

**Fig 5.**
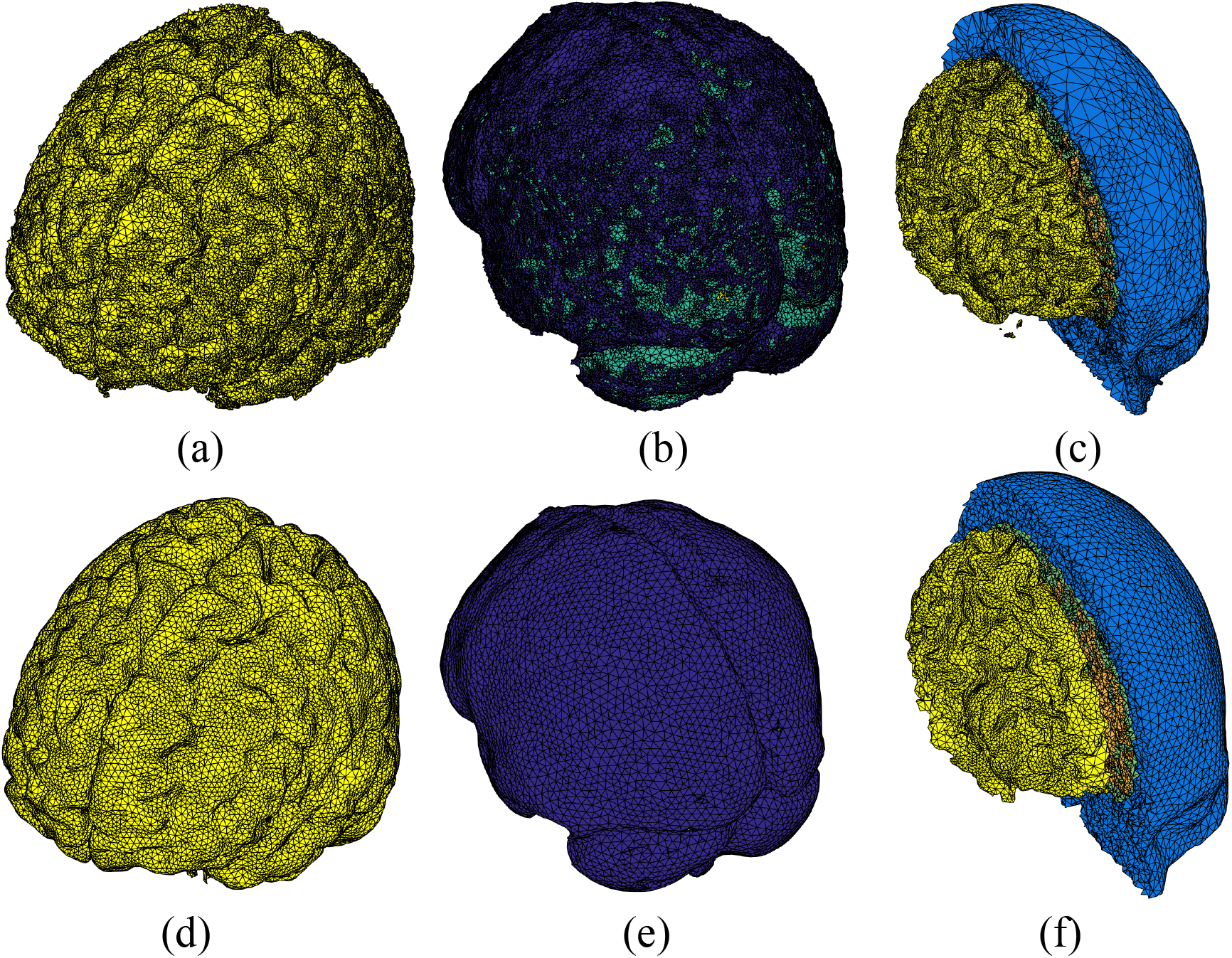
Comparison between (a-c) conventional CGAL-based volumetric meshing and (d-f) the new surface-based meshing approaches. From left to right, we show sample meshes for (a,d) GM, (b,e) CSF and (c,f) WM/scalp.

In Fig. 6, we show that users can generate tetrahedral meshes of different densities, by conveniently setting *V_max_* and the sizing-field (*S*) parameters. The set values are *V_max_* = 4, *S* =2 in Fig. 6(b), and *V_max_* = 2.5, *S* = 1.5 for Fig. 6(c). In the case of Fig. 6(c), we also reduced *R_max_* to 1.4 (in voxel length unit) for WM and GM, 1.7 for CSF, 2.0 for skull, and 2.5 for scalp.

**Fig 6.**
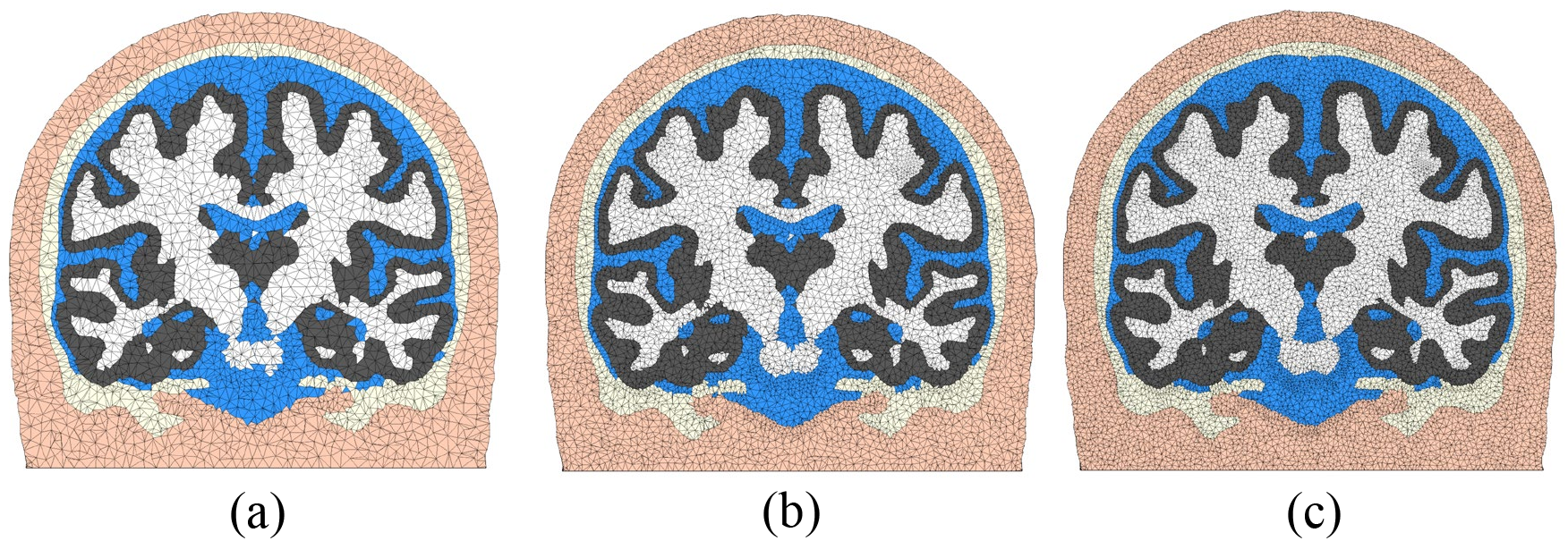
Demonstration of mesh density control. The mesh shown in (a) contains 181,026 nodes, 1,060,051 elements, with a runtime of 53.47 s. The mesh in (b) includes 499,134 nodes, 3,009,706 elements, with runtime 76.03 s, and (c) has 1,023,739 nodes, 6,220,187 elements and 135 s runtime. The 5 layers of brain tissues are, in order from outer-to-inner: scalp (apricot), skull (light-yellow), CSF (blue), GM (gray), and WM (white).

### 3.2 Hybrid meshing pipeline combining volumetric segmentations with tissue surface models

In this subsection, we demonstrate our “hybrid” meshing pathway (described in the right-half of Fig. 3). This approach combines tissue surfaces extracted from probabilistic segmentations with the pial and WM surfaces generated by dedicated neuroanatomical analysis tools, such as FreeSurfer and FSL. In the case of FreeSurfer, the raw pial and white matter surfaces are very dense. A mesh simplification algorithm is performed to “down-sample” the surface to the desired density. In Fig. 7, we characterize the trade-offs between mesh size and the surface error^54^ at various resampling ratios of the dense FreeSurfer-generated pial surface for the USC 30-34 atlas.^43^ The surface error is computed as the absolute value of the Euclidean distance (in mm) between each node of the down-sampled surface to the closest node on the original surface.^54^ From Fig. 7, at resampling ratio values below 0.1 (i.e. decimating over 90% edges), both gyri and sulci show a surface error above 0.5 mm. When the resampling ratio value is increased to 0.15, the observed error at the gyri becomes minimal; only a few regions show errors above 1 mm. A resampling ratio of 0.2 is selected in this example, giving a mean surface errors below 0.2 mm.

**Fig 7.**
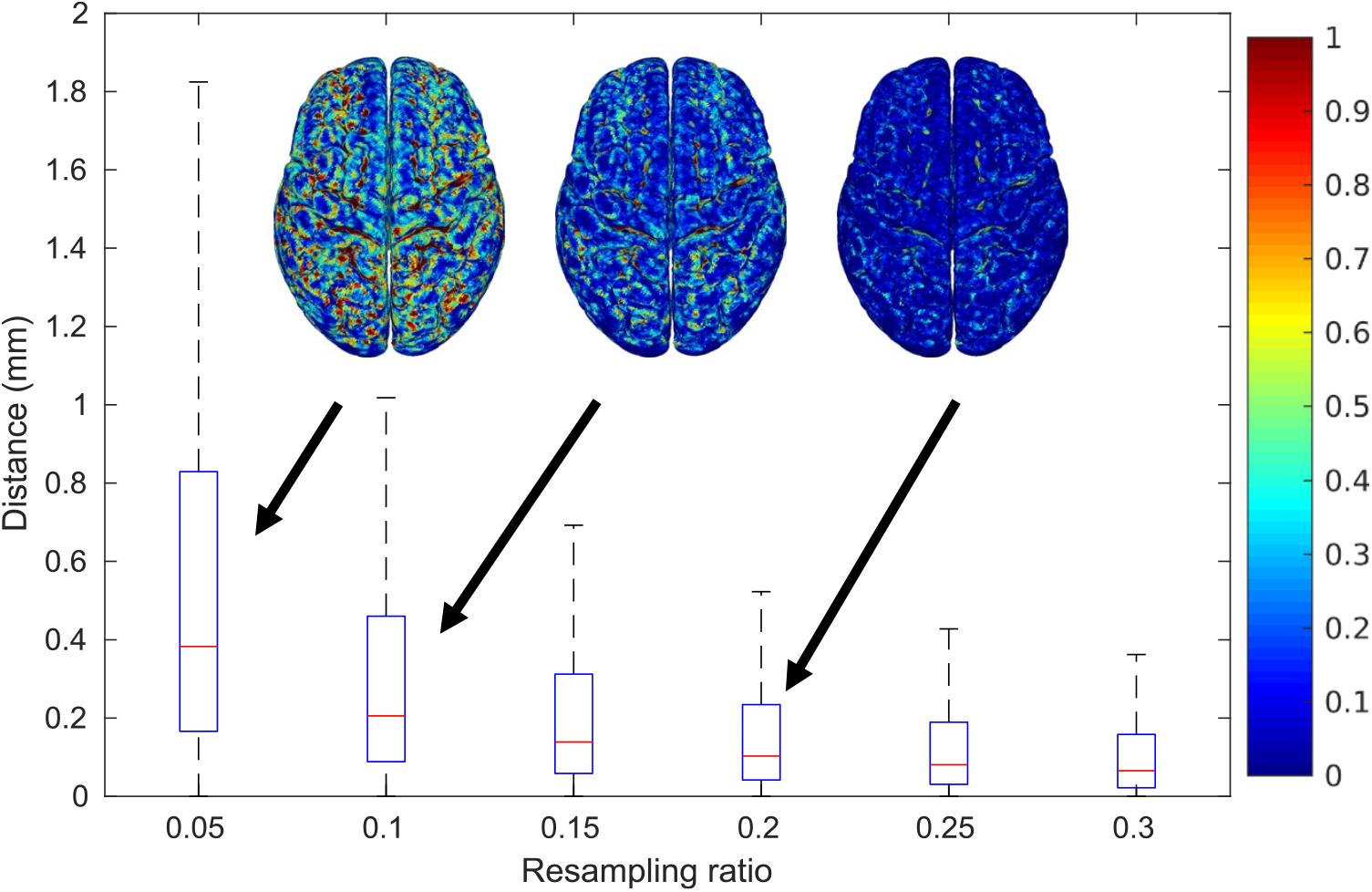
Box-plots of surface errors as a function of resampling ratio (percentage of edges that are preserved) when down-sampling a FreeSurfer-generated pial surface. Spatial distributions of the errors are shown as insets.

As we illustrate in Fig. 3, a number of additional step have been taken to process the FreeSurfer pial/WM surfaces. These include the merging of the left- and right-hemisphere surfaces, addition of the CSF ventricles, and the addition of cerebellum and brainstem using SPM segmentations. The final mesh, shown in Fig. 8, contains 150,999 nodes and 917,212 elements. The mesh generation time was 158.44 seconds. This meshing time is significantly lower than the reported 3-4 hours required for creating a similar mesh using “mri2mesh”.^35^ While not shown here, this hybrid workflow also accepts probabilistic segmentations produced by FSL or pial/WM surfaces created by BrainSuite and similar combinations.

**Fig 8.**
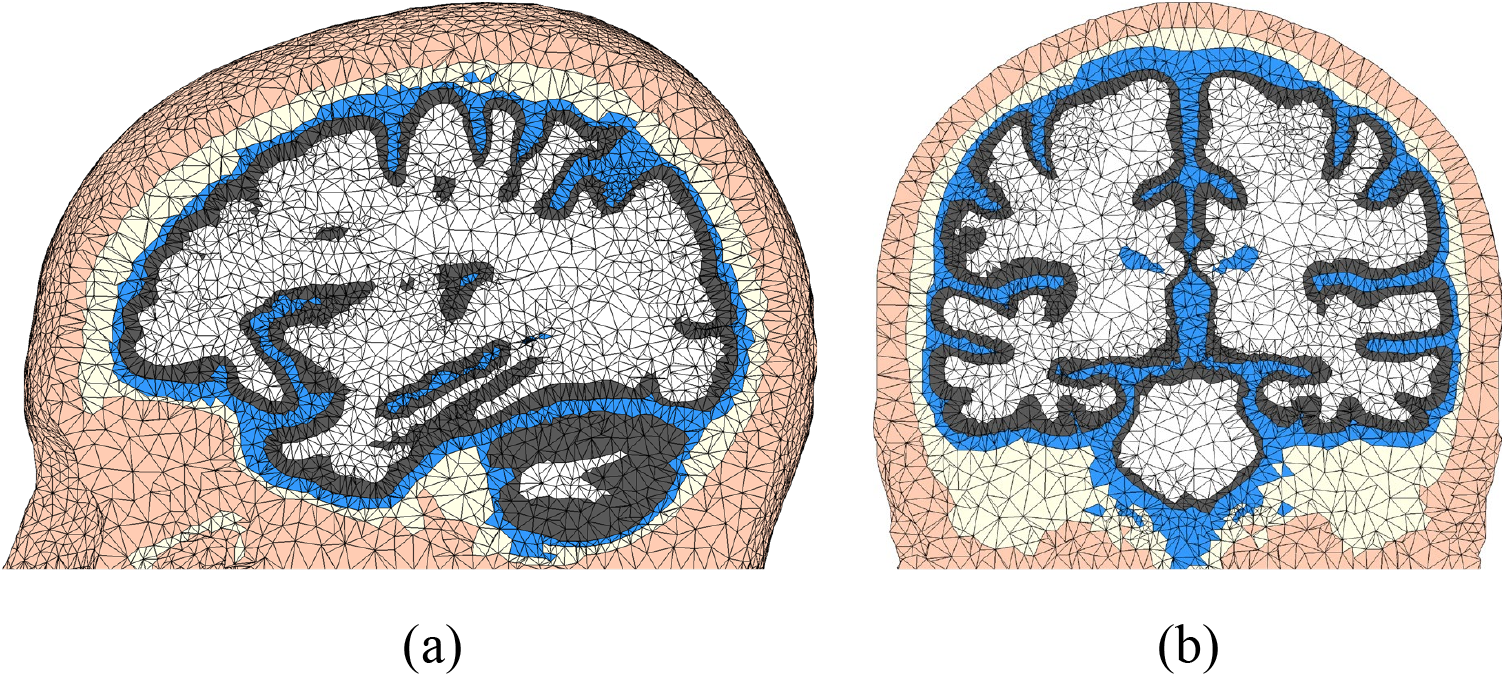
Tetrahedral mesh generated from a hybrid meshing pathway combining FreeSurfer surfaces with SPM segmentation outputs for the USC 30-34 atlas. The (a) sagittal and (b) coronal views are shown. The tissue layers include scalp (apricot), skull (light-yellow), CSF (blue), GM (gray), and WM (white).

### 3.3 Brain mesh library generated from public brain databases

To test the robustness of our meshing workflow described above, we successfully processed many of the publicly available brain segmentation datasets, including the BrainWeb database^55^ and the recently published Neurodevelopmental MRI database.^37^ For the BrainWeb atlas database, we created the corresponding mesh models directly from the available brain segmentations. For Neurodevelopmental atlases, the WM and GM segmentations provided as part of the the database were used; however, the CSF and bone segmentations were not directly included by the database because they are generally more difficult to create. For these missing tissues, separate segmentations for CSF and bone were created using SPM. In addition, the scalp surface was extracted from the raw MRI image using an intensity thresholding approach followed by 3 iterations of Laplacian+HC smoothing.^40^ In Fig. 9, we show 9 sample USC atlas brain meshes derived from adult and adolescent scans. In all processed MRI scans, the proposed meshing workflow worked smoothly; the average processing time is less than a minute per mesh when the voxel resolution is 1×1×1 mm^3^ and about 3 min per mesh when the resolution is 0.5×0.5×0.5 mm^3^.

**Fig 9.**
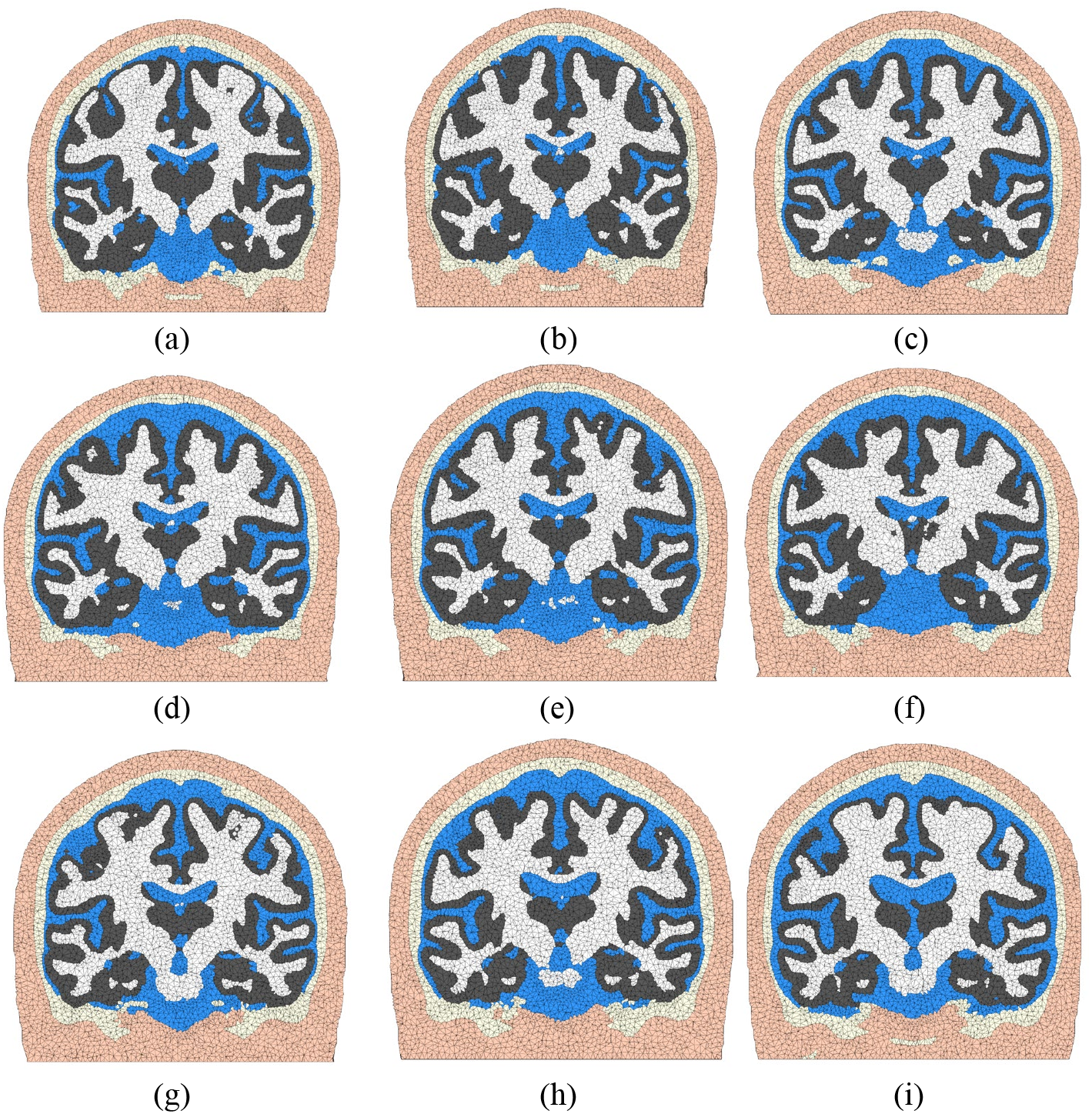
Illustrative brain mesh examples (coronal views) produced using the Neurodevelopmental MRI Database, including (a) 16 years, (b) 17.5 years, (c) 25-29 years, (d) 30-34 years, (e) 35-39 years, (f) 40-44 years, (g) 50-54 years, (h) 60-64 years, and (i) 70-74 years old. The tissue layers include scalp (apricot), skull (light-yellow), CSF (blue), GM (gray), and WM (white).

It is important to note that the CSF and bone segmentations in Fig. 9 have not been validated and are shown only for illustration purposes. In the initial release of our brain mesh library, we only included the GM/WM meshes from the Neurodevelopmental atlas database based on previously published segmentations.

### 3.4 Comparing mesh, voxel and layer-based brain models in light transport simulations

Next, we demonstrate the impact of different brain anatomical models, particularly between the mesh-, voxel- and layered-slab brain representations and highlight their discrepancies in optical parameters estimated from 3-D Monte Carlo light transport simulations. Here we use our in-house developed dual-grid mesh-based Monte Carlo (DMMC) simulator^17^ for mesh- and layer-based MC simulations, and MCX^31^ for voxel-based simulation. An MRI brain atlas (19.5 year group^51^) was selected for this comparison, although our methods are readily applicable to other brain models. The SPM segmentation (166×209×223 with 1×1×1 mm^3^ resolution) of the selected atlas and the generated tetrahedron-mesh from this segmentation are used for this comparison.

In this case, a tetrahedral mesh with 442,035 nodes and 2,596,064 elements is used for the DMMC simulations. The mesh was created using our aforementioned meshing pipeline with maximum-volume size *V_max_* = 30 mm^3^ and maximum Delaunay sphere radii *R_max_* = 1.2 mm for all tissue layers. In comparison, a much simplified layered-slab brain model is made of slabs of the same 5 tissue layers with the layer thicknesses calculated based on the mesh model: scalp: 7.25 mm, skull: 4.00 mm, CSF: 2.73 mm, and GM: 3.29 mm; WM tissue fills the remaining space. To minimize boundary effect, the layered-slab brain model has a dimension of 200×200×50 mm^3^.

For the two anatomically derived (voxel and mesh) brain simulations, an inward-pointing pencil beam source is placed at an EEG 10-5 landmark^56^ – “C4h” – selected using the “Mesh2EEG” toolbox.^57^ Within the same coronal plane, five 1.5 mm-radius detectors are placed on either side of the source along the scalp, 8.4, 20, 25, 30 and 35 mm from the source (in geodesic distance, see Fig. 10(a)), respectively, determined based on typical fNIRS system settings.^18, 58^ Similarly, for the simulations with the layered-slab brain model, a pencil beam source pointing down is placed at (99.5, 100, 0) mm. A similar set of detectors are placed on one side of the source due to symmetry, see Fig. 10(b).

**Fig 10.**
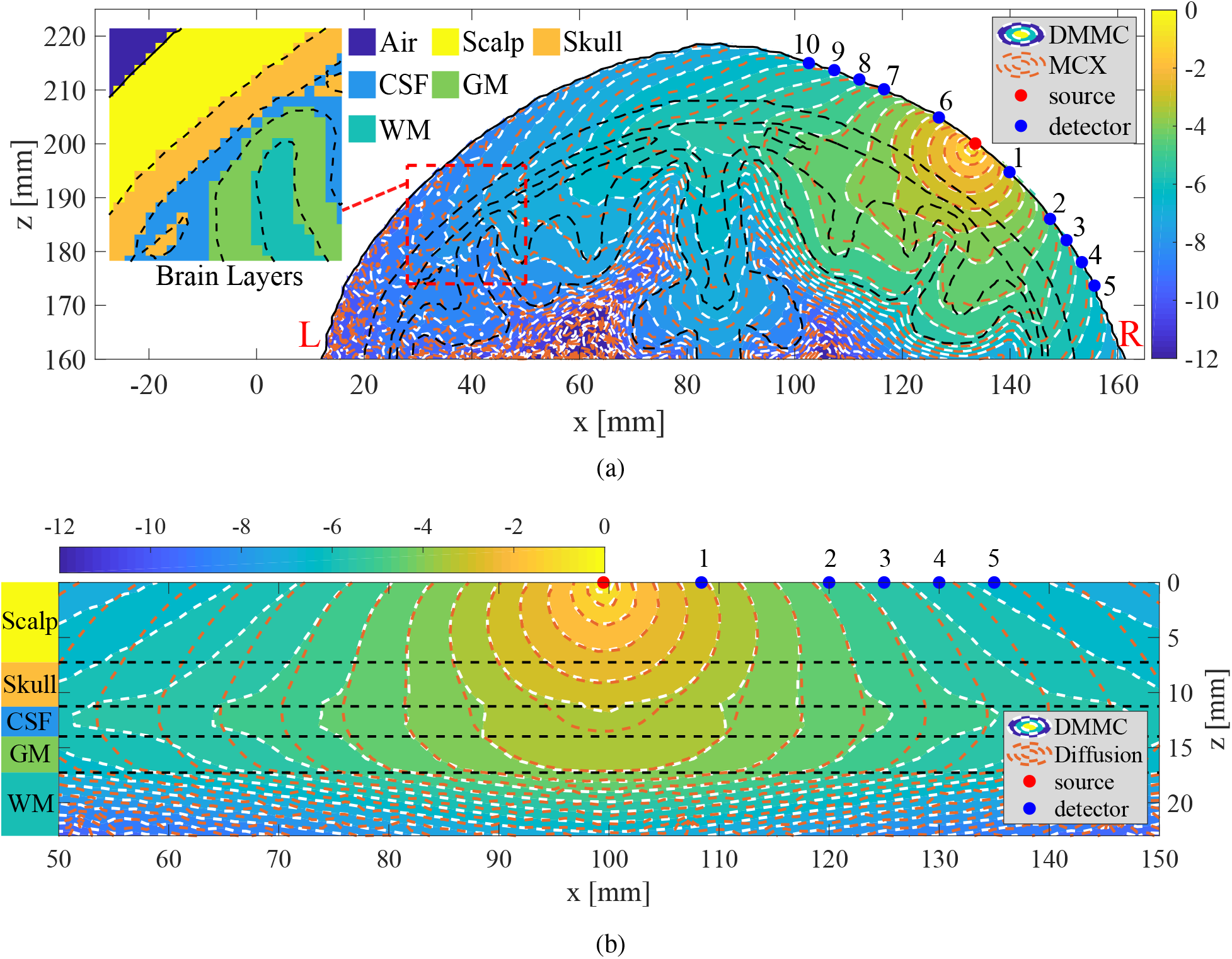
Comparisons of fluence distributions in an MRI brain atlas (19.5 year) using 3 different brain models: (a) MC fluence maps using anatomically derived mesh (computed using DMMC) and voxel (computed using MCX) brain representations, and (b) fluence maps computed using the MC and diffusion approximation in a simple layered-slab brain model. Contour plots, in log-10 scale, are shown along the coronal planes with each brain tissue layer labeled and delineated by black dashed-lines. In (a), the “L” and “R” markings (red) indicate the left and the right brain, respectively. The comparisons between the mesh and voxel tissue boundaries are shown in the inset of (a).

Three-dimensional MC photon simulations are performed on all 3 brain models, where DMMC is used for both the mesh-based and the layered-slab brain models, and MCX is used for the voxel-based brain model. The output fluence distributions along the source-detector plane are compared in Fig. 10(a). From these MC simulations, we also report the average partial-pathlengths in the brain regions (*PPL_B_*), average total-pathlengths (*TPL*) and their percentage ratios 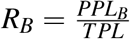 in Table 1. Moreover, we also computed the percentage fraction of the energy deposition in the gray matter region with respect to the total simulated energy. In addition to MC-based photon modeling, we have also applied the diffusion approximation, frequently seen in the literature,^5^ to the layered-slab brain model and compare the results with those derived from the MC method in Fig. 10(b). For solving the diffusion equation, we used our in-house diffusion solver, Redbird.^59^ The reduced scattering coefficient of the CSF region is set to 0.3 mm^−1^ as suggested in Refs.^5, 48^ The Redbird solution matches excellently with that from NIRFAST^15^ (not shown).

**Table 1.** Comparison of key optical parameters derived from MC simulations from an MRI brain atlas (19.5 year): For each detector, we compare the average photon partial-pathlengths in the brain region (*PPL_B_*), total-pathlengths (*TPL*) and their percentage ratios (*R_B_*) derived from mesh-based (DMMC), voxel-based (MCX) and layered-slab (DMMC) brain representations at various source-detector separations (SD).

|  |  | Mesh-based brain model (DMMC) |  |  | Voxel-based brain model (MCX) |  |  | Layered-slab brain model (DMMC) |  |  |
| --- | --- | --- | --- | --- | --- | --- | --- | --- | --- | --- |
| det. # | SD (mm) | PPL<sub>B</sub> (mm) | TPL (mm) | R<sub>B</sub> | PPL<sub>B</sub> (mm) | TPL (mm) | R<sub>B</sub> | PPL<sub>B</sub> (mm) | TPL (mm) | R<sub>B</sub> |
| 1 | 8.4 | 0.05 | 34.37 | 0.14% | 0.06 | 33.91 | 0.17% | 0.13 | 36.52 | 0.35% |
| 2 | 20 | 1.55 | 92.96 | 1.66% | 1.90 | 93.53 | 2.03% | 3.03 | 101.9 | 2.97% |
| 3 | 25 | 4.33 | 122.4 | 3.54% | 5.33 | 123.6 | 4.31% | 7.05 | 133.4 | 5.28% |
| 4 | 30 | 8.74 | 150.1 | 5.82% | 9.73 | 150.1 | 6.48% | 12.5 | 163.5 | 7.64% |
| 5 | 35 | 12.8 | 169.6 | 7.56% | 14.4 | 170.5 | 8.44% | 17.4 | 186.5 | 9.34% |
| 6 | 8.4 | 0.04 | 36.06 | 0.11% | 0.05 | 36.10 | 0.14% |  |  |  |
| 7 | 20 | 1.20 | 95.47 | 1.26% | 1.38 | 97.16 | 1.42% |  |  |  |
| 8 | 25 | 3.48 | 125.5 | 2.78% | 3.70 | 126.4 | 2.92% |  |  |  |
| 9 | 30 | 7.77 | 157.0 | 4.95% | 7.90 | 158.0 | 5.00% |  |  |  |
| 10 | 35 | 12.2 | 179.5 | 6.78% | 12.0 | 180.1 | 6.68% |  |  |  |

In Fig. 10(a), the fluence contour plots produced by MCX (orange dashed-line) and DMMC (white dashed-line) agree excellently in the vicinity of the source, while noticeable discrepancies are observed when moving away from the source. We believe this is a combined result of 1) photon energy deposition variations due to the small disagreement between a terraced tissue boundary and the smooth surface boundary, and 2) the distinct photon reflection behaviors between a voxel- and a mesh-based surface due to the differences in the orientations of surface facets. The effect from the first cause is largely depicted by the deviation between the two solutions in the depth direction near the source. Such difference is particularly prominent near highly-curved boundaries or near boundaries with high absorption/scattering contrasts, such as the CSF region beneath detectors #7, #8 and #9 in this plot. The effect from the second cause is highlighted by the worsened discrepancy when moving away from the source along the scalp layer, for example, the scalp region to the left of detector #10. Overall, the second source of error is noticeably prominent than the mismatch resulted from the first cause. This observation is further validated by disabling the refractive-index mismatch calculations in both of our simulations (results not shown): the error along the scalp surface was largely removed, but the deviations in the deep-brain regions remain.

In Fig. 10(b), the diffusion approximation (orange dashed-line) and MC (white dashed-line) produce well-matched fluence contour plots near the source but show significant difference in the regions distal to the source. The difference is particularly noticeable within the CSF and GM regions and above 30 mm source-detector separation on the scalp surface. We believe it is largely due to the error introduced by the approximated CSF reduced scattering coefficient.^48^

To further quantify the differences caused by different brain representations and their impact to fNIRS brain measurements, in Table. 1, we also compare several key photon parameters derived from MC simulations. Here we use the parameters derived from mesh-based MC as the reference. This is because MC solutions are typically used as gold-standard, and mesh-based shape representations are known to be more accurate than voxelated domains.^16^ We observe that simulations using voxel-based and layered-slab brain models tend to overestimate *PPL_B_* compared to the mesh models. For detectors #6 to #10, the voxel-based simulation gives a 21% over-estimation at the shortest source-detector (SD) separation (8.4 mm); such discrepancy is reduced to within ±2% at the largest two separations (30 and 35 mm). The *TPL* values are less susceptible to anatomical model accuracy, reporting a percentage difference between 0.1% to 1.8%. As a result, the difference in *R_B_* is largely modulated by that of *PPL_B_*, ranging between −1.5% to 21%. However, for detectors #1 to #5 located on a different brain region where the superficial layers are shallower than those under detectors #6 to #10, more pronounced over-estimations of *PPL_B_* for all SD separations, ranging from 12% to 23%, are observed, resulting in an *R_B_* percentage difference between 12% and 24%. Similarly, compared to mesh-based model, the layered-slab brain MC simulations report significant over-estimation of *PPL_B_* (36% to 166% with the highest difference at the shortest separation) and *TPL* (6.3% to 10%), resulting in significant variation in *R_B_*: 151% at the shortest SD separation and 78% to 24% for four long separations. Furthermore, we have also computed the percentage fraction of the energy deposition within the GM. This fraction is 1.69% when using mesh-based brain model, and 1.42% when using a voxel-based brain model, resulting in a 16% reduction in brain energy deposition. This result could have some implications to many photobiomodulation (PBM) applications.^50^

## 4 Conclusion

In this work, we address the increasing needs for accurate and high-quality brain/head anatomical models that arise in fNIRS and many other neuroimaging modalities for brain function quantification, image reconstruction, multi-physics modeling and visualization. Combined with the advance in light transport simulators,^16, 17, 47^ our proposed brain mesh generation pipeline enables fNIRS research community to utilize more accurate anatomical representations of the human brain to improve quantification accuracy, and make atlas-based as well as subject-specific fNIRS analysis more feasible. This also gives us an opportunity to systematically investigate how neuroanatomical models – ranging from the simple layered-slab brain model to voxel-based and mesh-based models – impact the estimations of optical parameters that are essential to fNIRS imaging.

Specifically, we first described a fast and robust brain mesh generation algorithm and demonstrated that our MATLAB-based open-source toolbox, “Brain2Mesh”, can produce high-quality brain and full-head tetrahedral meshes from multi-label or probabilistic segmentations with full automation. The abilities to create tissue boundaries from gray-scale probabilistic maps and incorporate detailed surface models from FreeSurfer/FSL ensure smoothness and high accuracy in representing brain tissue boundaries. The output meshes generally exhibit excellent shape quality without needing to generate excessive number of small elements, such as in many existing mesh generators. For most of the included examples, the processing time ranges between 1 and a few minutes using only a single CPU thread. This is dramatically faster than most previously published brain meshing tools.^30, 35, 60^ Moreover, the entire meshing pipeline was developed based on our open-source meshing toolbox, Iso2Mesh, and other open-source meshing utilities such as CGAL, TetGen and Cork. This ensures excellent accessibility of this tool to the community. In addition to developing this brain mesh generation toolkit, we have also produced a set of high-quality brain atlas mesh models, including the widely used BrainWeb, Colin27 and MNC atlases. We believe these ready-to-use brain/fullhead models will be valuable resources for the fNIRS community.

Another important aspect of this study is that we demonstrate how tissue boundary representations, especially layered-, voxel- and mesh-based anatomical models, could impact light transport simulations in fNIRS data analysis. While the modeling error caused by voxelization in MC simulations has been previously reported,^61^ we believe this is the first time such discrepancy has been quantified, particularly in the context of brain imaging, enabled by our unique access to high-quality brain meshes and highly accurate mesh-based MC simulation tools. We believe such findings could provide guidance for advancing fNIRS towards improved accuracy and broad utility. Our open-source meshing software as well as the brain mesh library are freely available at http://mcx.space/brain2mesh.

## Acknowledgments

The authors wish to thank the funding supports from the National Institutes of Health under grant numbers R01-GM114365, R01-CA0204443 and R01-EB026998. We also acknowledge the valuable inputs from Dr. Hang Si on the use of TetGen, as well as the instructive conversation with Dr. John Richards and Dr. Katherine Perdue on brain segmentation.

